# Pelleted-hay alfalfa feed increases sheep wether weight gain and rumen bacterial richness over loose-hay alfalfa feed

**DOI:** 10.1101/605113

**Authors:** Suzanne L. Ishaq, Medora M. Lachman, Benjamin A. Wenner, Amy Baeza, Molly Butler, Emily Gates, Sarah Olivo, Julie Buono Geddes, Patrick Hatfield, Carl J. Yeoman

**Affiliations:** Biology and the Built Environment Center, University of Oregon, Eugene, OR; Department of Animal and Range Sciences, Montana State University, Bozeman, MT; Department of Animal Sciences, The Ohio State University, Columbus, OH

**Keywords:** bacteriome, diet-microbe interactions, feed digestibility, particle size

## Abstract

Diet composed of smaller particles can improve feed intake, digestibility, and animal growth or health, but in ruminant species can reduce rumination and buffering – the loss of which may inhibit fermentation and digestibility. However, the explicit effect of particle size on the rumen microbiota remains untested, despite their crucial role in digestion. We evaluated the effects of reduced particle size on rumen microbiota by feeding long-stem (loose) alfalfa hay compared to a ground and pelleted version of the same alfalfa in yearling sheep wethers. *In situ* digestibility of the pelleted diet was greater at 48 h compared with loose hay; however, distribution of residual fecal particle sizes in sheep did not differ between the dietary treatments at any time point. Both average daily gain and feed efficiency were greater for the wethers consuming the pelleted diet. Observed bacterial richness was very low at the end of the adaptation period and increased over the course of the study, suggesting the rumen bacterial community was still in flux after two weeks of adaptation. The pelleted-hay diet group had a greater increase in bacterial richness, including common fibrolytic rumen inhabitants. The pelleted diet was positively associated with several *Succiniclasticum*, a Prevotella, and uncultured taxa in the Ruminococcaceae and Rickenellaceae families and Bacteroidales order. Pelleting an alfalfa hay diet for sheep does shift the rumen microbiome, though the interplay of diet particle size, retention and GI transit time, microbial fermentative and hydrolytic activity, and host growth or health is still largely unexplored.

## Introduction

It has been well established that the nutrient composition of a diet affects the gastrointestinal tract (GIT) microbiota [1,2], yet the physical structure and complexity of the diet may also alter its interactions with the GIT microbiota and host. In cattle, longer fiber particles have been shown to improve rumination [3] and ultimately fiber digestibility [4]. However, shorter or smaller diet particles can change the dynamics of digestion and produce a number of favorable outcomes. Specifically, reductions in particle size associated with mastication or mechanized processing correspond to increases in feed surface area and thus allow for greater microbial attachment and relative fibrolytic and fermentative activities as has been shown *in vitro* [5]. Mechanical breakdown during diet preparation can also physically disrupt waxy plant cuticles and cell walls that can otherwise impede microbial attachment and degradation, thus making plant carbohydrates more available [6] and decreasing the potential confounding effect of forage fragility within the rumen [7]. Hydrolytic activities are also likely further enhanced by the reductions in buoyancy and increased functional specific gravity [8] associated with reduced particle size, that would allow smaller particles to sink beneath the dorsally-located rumen mat and into the microbe-rich rumen liquor [9,10].

Particle size not only affects the amount of surface area available for microbial attachment [11], but also the retention time of feed within the rumen environment. Feed has been shown to remain confined to the rumen until it reaches a critical size threshold – demonstrated by a number of studies to range from ∼1 to 1.18 mm [12–14] after which it can escape the filtering capacity of the omasum [3,15–17]. Consequently, reducing the particle size of ruminant diets has been seen to increase GIT passage rate [18,19] and can thereby free up additional capacity for rumen fill which encourages dry matter intake (DMI). Both increased DMI and increases in hydrogen gas partial pressure that can result from rapid fermentation can further increase GIT passage rate and reduce ruminal retention time independent of particle size [12–14]. After exiting the rumen via the omasum, digesta proceeds to the highly acidic abomasum wherein microbial activity is largely halted [20]. Thus, the increased potential for fibrolytic activity associated with reductions in feed particle size are offset by the shorter time available for these activities to occur. Within this context, the effect of feed particle size on rumen microbial fibrolytic activity remains largely unexplored.

Additional factors that can further complicate the relationship of feed particle size and digestibility include the knowledge that ingesting smaller particles reduces the amount of time spent chewing [15,21–23] and ruminating [3,16,17]. Further, less mastication and rumination decreases the amount of the buffered saliva being swallowed [21,24], and as saliva is responsible for neutralizing 30 to 50% of the acid resulting from fermentation [25,26] this can lead to acidification of the rumen [25], and associated reductions in microbial attachment and fibrolytic activity [27–29]. Bacterial ionic charge is facilitated by cell walls composition; negatively-charged bacteria are better able to attach to lignocellulosic material, often mediated through binding to positively-charged magnesium [30]. The downstream impact on animal performance is further complicated. The solubility of dietary nutrients are also altered by the physics of particle size, thus affecting their ease of host digestion, including transit through mucus, and absorption by epithelia [11,31]. Similarly, small or homogenous particle sizes have been reported to preclude normal papillae development [32,33], although these alterations have been either reported to be localized within the GIT [33] or not significantly altered [34–36]. Other studies reported that roughage decreased omasal size and epithelial keratinization [34], and increased rumen wall muscularity over starch-based diets [35,36]. Lignin, cellulose, and hemicellulose can form superstructures in the luminal digesta, making digestible carbohydrates less accessible, and further contributing to nutrient loss through the binding of metal ions [30]. Thus, decreasing particle size has been demonstrated to affect the lignin content of feed by physical disruption [22], and can improve access to fiber and microbial fermentation *in vitro* and *in vivo* [22,23,37–39]. Overall the influence of particle size on animal performance, measured as feed efficiency or weight gain, is unclear with several studies having reported improvements with decreased particle size [23,32,40] while others do not [41–43]. These differences are likely the result of varied rumen conditions imposed by reduced particle size or other confounding conditions.

The objective of this research was to determine the effect of particle size reduction on animal growth, feed intake, feed digestibility, and microbial diversity by comparing lambs fed either a long-stem second-cutting alfalfa hay or the same alfalfa after pelleting. Our working hypothesis was that decreasing particle size of the alfalfa hay and delivery in the form of a pellet would increase animal growth and feed digestibility due to reduced particle size. Further, pelleted feed was hypothesized to decrease microbial diversity by allowing for more rapid fermentation and alteration of environmental conditions in the rumen.

## Materials and Methods

### Ethics Statement

All animal procedures were approved by the Montana State University Agricultural Animal Care and Use Committee (Protocol #2014-AA04). The trial was conducted at the Montana State University Bozeman Agriculture Research and Teaching Farm in Bozeman, MT, USA.

### Diet

Second cutting alfalfa was harvested and hayed in fall 2014 and used to produce 2 dietary treatments. The first treatment consisted of the loose, long-stem alfalfa hay, and the second treatment comprised the same hay being commercially ground and pelleted. Nutritional profiles of each diet were determined via NIR by Dairy One Forage Lab (Ithaca, NY) and are provided in Table 1. While it was intended to have the diets balanced for NDF, ADF, CP, and ME, the mechanical grinding and pelleting process induced some unavoidable changes to the forage provided that resulted in minor differences between diets among some nutrient categories. Since the pellets were derived from the same hay source, the primary contributing factors to variation between treatments was likely shatter of leafy particles from hay at grinding and pelleting, and the addition of some oil to properly bind the pellet. Fiber (NDF and ADF) were decreased by the pelleted diet while CP was increased.

**Table 1:** Nutritional composition of loose-hay and pelleted-hay alfalfa diets, as well as *in situ* rumen digestibility.

|  | Loose feed | Loose refusals | Pelleted feed | Pellet refusals |
| --- | --- | --- | --- | --- |
| Moisture | 8.6 | 7.9 | 10.2 | 9.4 |
| Dry Matter (DM; %) | 91.4 | 92.1 | 89.8 | 90.6 |
| Crude Protein (CP; % DM) | 16 | 13 | 20 | 20 |
| Non-Fiber Carbohydrates (NFC) | 18.2 | 16.8 | 22.5 | 23.2 |
| Neutral-Detergent Fiber (NDF; % DM) | 54.6 | 59.7 | 46.8 | 46.2 |
| peNDF<sub>1.18</sub> (% DM) | 44.3 |  | 14.4 |  |
| Acid-Detergent Fiber (ADF; % DM) | 43.8 | 49.1 | 37.9 | 37.5 |
| Lignin (% DM) | 10.8 | 11.8 | 9.3 | 9 |
| Ash (% DM) | 6.9 | 5.8 | 10.4 | 9.9 |
| Total Digestible Nutrients (TDN) | 52 | 49 | 58 | 58 |
| Relative Feed Value (RFV) | 93 | 79 | 118 | 120 |
| % Digested 24 hr *in situ* ($P = ns$) | 50.36% | | 56.30% | |
| % Digested 48 hr *in situ* ($P < 0.001$) | 51.50% | | 58.80% | |

### Experimental Design

Rambouillet wethers (n = 10, avg. BW = 30 ± 5 kg) were randomly assigned to either loose-hay or pelleted diets within blocks by weight. Animals were adapted to a 50:50 mix of both dietary treatments over 14 d prior to the start of the trial, with the final 7 d being spent in metabolism crates with fastened fecal bags for acclimation. The end of the adaptation period was considered time point zero (day 0/week 0), following which the animals were fed their assigned diet for a 14-day period. Wethers were fed *ad libitum* twice daily between the times of 0700 to 0830 and 1730 to 1900. Orts and feces were collected; water was available free choice at all times. Animals were weighed once weekly, and blood and rumen samples collected at the start of the trial (day 0, end of adaptation period) and at days 7 and 14. Rumen samples were collected using sterilized foal tubes with a 600cc large animal dosing syringes, and immediately stored at −20°C until processing (within 3 weeks). Feed samples were randomly taken in triplicate daily from hay bales using a Penn State Forage Sampler (loose) or pellet bags (pellet) and pooled by treatment day to submit a single feed sample for laboratory analysis. Following weighing, feed samples and orts were stored in paper bags at ambient temperature until grinding. Feces were collected and frozen at −20°C to prevent microbial metabolism.

### Intake and digestibility

Intake was determined as feed provided less orts. Dry matter (DM) of feed, orts, and feces was determined by drying overnight in an oven set to 100 °C. Digestibility of alfalfa treatments was estimated by 48 h *in situ* (i.e. *in sacco*) incubation in a rumen-cannulated dairy cow using a standardized procedure as previously described [44]. Particle size distributions of feed, orts, and feces were determined by wet sieving as previously described [45] using a set of 12 mesh sieves (Sigma Aldrich, St. Louis, MO, USA) ranging from No. 4 (> 4.76mm) down to No. 200 (0.074 - 0.087 mm) Using lignin as an internal marker, fecal nutrient composition was used to predict apparent nutrient digestibility (DM, CP, NDF, ADF) using the equation [46]:

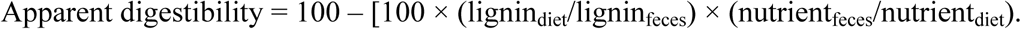

Means for sheep intake and digestibility were compared in a mixed effects model (SAS 9.4, Cary, NC) with fixed effect of treatment, random effect of sheep, and a repeated sampling day. Repeated covariance structure was selected using the lowest BIC and significance was declared at *P* < 0.05. Means for sheep gain and feed conversion ratio (kg gain/ kg DMI) used the same structure with the removal of repeated measures since there were only 3 weekly samples included. Means for *in situ* digestibility were compared via linear mixed modeling (lmer) and least squares means (lsmeans [47]), with sheep as a fixed effect to control for repeated measures, and significance was designated at *P* < 0.05 with tendencies from 0.05 *≤ P* < 0.10.

### Blood panel

Blood samples (10 mL) were collected from each wether by jugular venipuncture. All samples were collected using serum vacutainers (Becton Dickinson, Franklin Lakes, NJ) fitted with 20-gauge vacutainer needles. Serum tubes were allowed to coagulate before being placed on ice and transported to the Montana Veterinary Diagnostic Laboratory (Bozeman, MT, USA) where large animal biochemistry panels were performed using an automated biochemical analyzer. The panel included creatine kinase/phosphokinase (CPK), creatinine (CREAT), aspartate aminotransferase (AST), alkaline phosphatase (ALKP), total protein (TP), albumin (ALB), globulin (GLOB), glucose (GLU), blood urea nitrogen (BUN), calcium (Ca), phosphorus (P), sodium (Na), potassium (K), chloride (Cl), magnesium (Mg), total bilirubin (T Bili), direct bilirubin (D Bili), and total carbon dioxide (TCO2). Means for blood measurements (Supplementary Table 1) were compared in a mixed effects model (SAS 9.4, Cary, NC) with fixed effect of treatment, random effect of sheep, and a repeated sampling day. Repeated covariance structure was selected using the lowest BIC and significance was declared at *P* < 0.05 with tendencies from 0.05 *≤ P* < 0.10.

**Supplemental Table 1.**
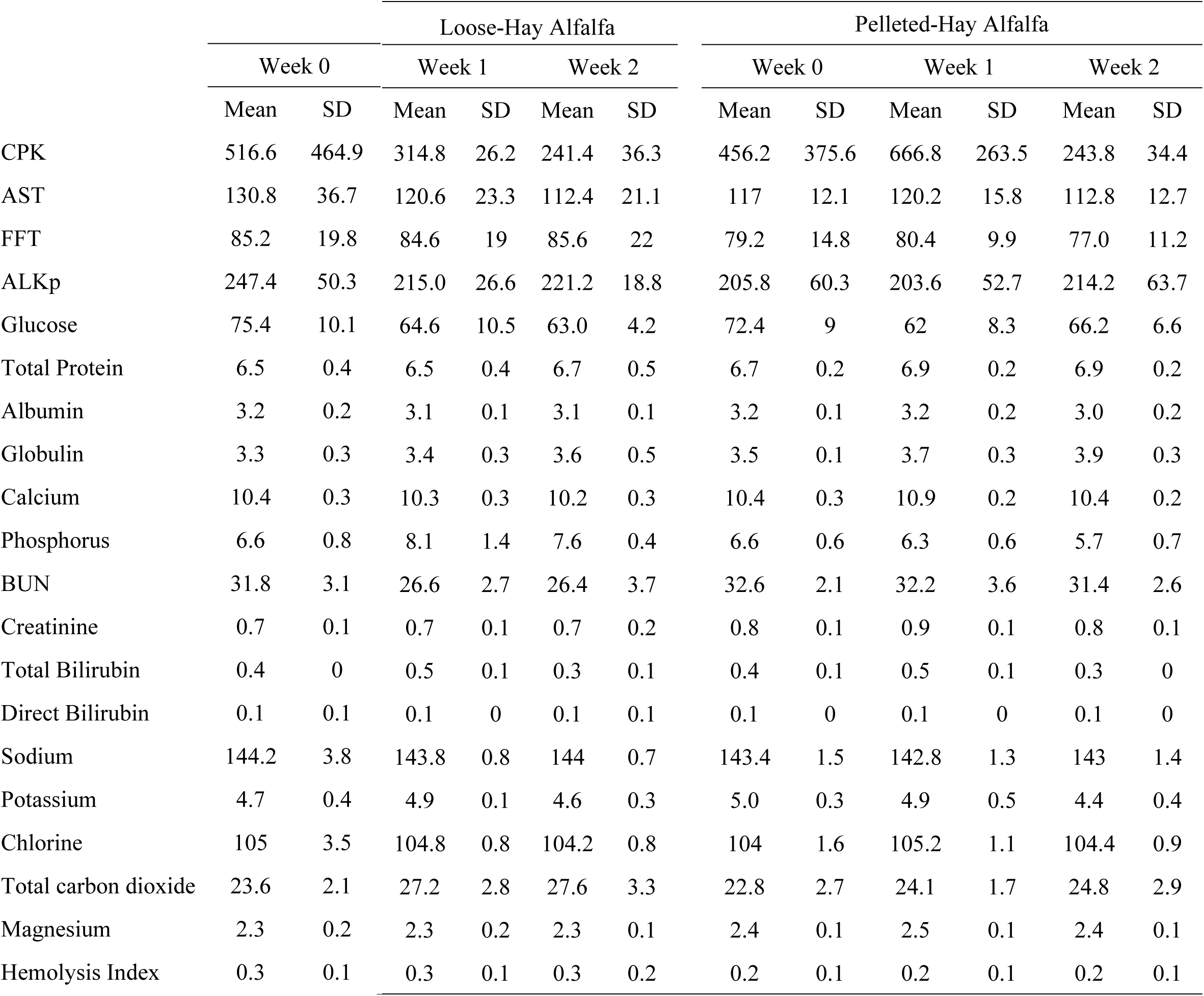
Serum parameters for wethers receiving either a loose-hay or a pelleted-hay alfalfa diet treatment for two weeks. Icteric index and Lipemic index were both 0.17 for all wethers.

### Bacterial V3-V4 16S rRNA Profiles

Rumen samples were thawed on ice, and bulk nucleic acids extracted via the MoBio PowerFecal DNA Isolation kit (Carlsbad, CA, USA) following manufacturer protocols. PCR amplification, amplicon quantification and purification, and Illumina MiSeq sequencing were performed as previously described [48] using the V3-V4 region of the 16S rRNA gene using a 300-cycle kit. Forward sequences were processed using the DADA2 package [49] in R [50], using filterAndTrim command parameters: trimLeft = 10, truncLen=270, maxN=0, maxEE=2, truncQ=2, rm.phix=TRUE. Sequences had taxonomy assigned with the Silva ver. 132 taxonomy database, sequences identified as chloroplast or mitochondrial were removed, as were chimeras. Negative controls included at DNA extraction through to sequencing were used to remove sequences from samples which also appeared in negative controls, as per (https://github.com/SueIshaq/Examples-DADA2-Phyloseq). Workflow was validated using an in-house 32 member mock community, as previously described [51]. Per sample, the number of sequences which passed quality-control measures ranged from 3,520 – 221,441, and for comparison, SVs were subsampled down to 3,520 per sample. Remaining sequences were analyzed using packages phyloseq [52], DESeq [53], rfPermute [54], and visualized with ggplot2 [55]. Observed richness values were determined to not be normally distributed via a Shapiro-Wilks Test, and were analyzed using a generalized linear mixed-effects model (GLMM) with lme4 [56], using Sheep ID as a fixed effect. Importance variables were identified using distance-based redundancy analysis (dbRDA; capscale) and variance partitioning (varpart). Community distance was assessed using unweighted Jaccard distance and adonis (permanova) in vegan [57], with Sheep ID as a fixed effect. Significance was designated at *p* < 0.05. Raw sequence reads are available from NCBI SRA under BioProject Accession number PRJNA384590.

## Results

### Intake, particle size, digestibility

The two diets differed in their particle size distributions (Fig. 1). Loose-hay alfalfa stems were an average 109.8 ± 44.9 mm in length, and 59.7% of hay particles were retained above the 4.76 mm screen (largest sieve used) while the majority (57.7%) of pelleted alfalfa feed was between 0.42 and 2.37 mm. Importantly, 18.4% and 69.2% of mass of the two diets, respectively, were less than the 1.18 mm threshold previously reported [12–14] to be the critical size for feed to escape the rumen. Calculated peNDF_1.18_ [58] of the loose hay and pelleted diets were 44.3% and 14.4%, respectively. Particles from hay refusals (orts) averaged 95.6 mm ± 60.1 in length, thus, we concluded that sorting against large particles did not occur in the loose hay diet. Following consumption and digestion, distribution of residual fecal particle sizes did not differ between diets at any time point sampled (Fig. 1, Supp. Table 2; *P* > 0.05).

**Figure 1.**
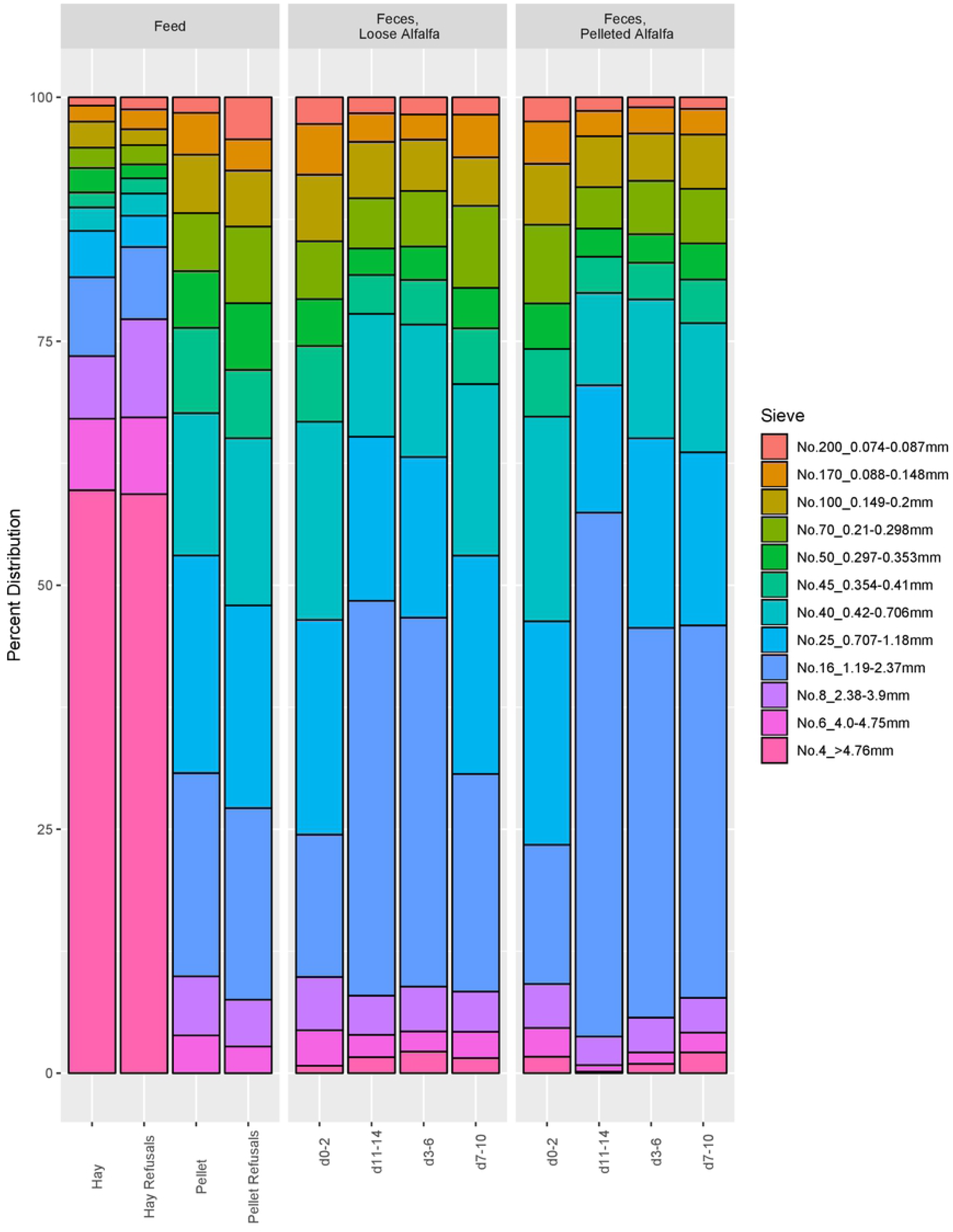
Particle size distribution of in diet treatment biomass, and percentage of residual feed particles in feces in wethers on a loose-hay (Hay) and pelleted (Pellet) alfalfa diet.

**Supplemental Table 2.** Residual nutritional composition of feces from wethers on loose-hay or pelleted-hay alfalfa diets over the two-week experimental period. Crude protein (% DM) was significantly different by diet (lmer, *p* = 0.00150) and day (lmer, *p* = 0.00285) overall.

| Day | Moisture |  | DM (%) |  | CP (%DM) |  | ADF (%DM) |  | NDF (%DM) |  | Lignin |  |
| --- | --- | --- | --- | --- | --- | --- | --- | --- | --- | --- | --- | --- |
|  | Mean | SD | Mean | SD | Mean | SD | Mean | SD | Mean | SD | Mean | SD |
| **Loose-Hay Alfalfa** |  |  |  |  |  |  |  |  |  |  |  |  |
| **0-2** | 1.7 | 0.4 | 98.3 | 0.3 | 15.5 | 0.8 | 44.3 | 5.7 | 57.6 | 3.4 | 17.5 | 2.5 |
| **3-6** | 1.5 | 0.4 | 98.5 | 0.4 | 16.6 | 0.7 | 41.2 | 2.9 | 53.3 | 3.2 | 17.1 | 1.2 |
| **7-10** | 1.6 | 0.7 | 98.4 | 0.7 | 15.3 | 0.3 | 42.2 | 5.0 | 54.8 | 3.5 | 16.9 | 2.6 |
| **11-14** | 1.4 | 0.3 | 98.7 | 0.3 | 14.8 | 0.8 | 41.4 | 2.9 | 55.8 | 1.4 | 16.4 | 1.3 |
| **Pelleted-Hay Alfalfa** |  |  |  |  |  |  |  |  |  |  |  |  |
| **0-2** | 1.4 | 0.8 | 98.6 | 0.8 | 14.7 | 0.6 | 43.5 | 3.7 | 58.4 | 2.5 | 14.8 | 1.2 |
| **3-6** | 1.0 | 0.5 | 99.0 | 0.5 | 14.3 | 0.5 | 45.8 | 3.0 | 59.5 | 2.0 | 16.0 | 1.9 |
| **7-10** | 1.0 | 0.4 | 99.0 | 0.4 | 14.4 | 0.4 | 44.2 | 1.9 | 55.4 | 0.8 | 15.9 | 1.3 |
| **11-14** | 0.9 | 0.4 | 99.1 | 0.3 | 13.8 | 0.7 | 44.1 | 2.7 | 58.2 | 3.9 | 15.9 | 1.7 |

Nutritional analyses of the two diet treatments are shown in Table 1. *In situ* dry matter disappearance at 48 h indicated that the pelleted diet was more digestible than the whole hay diet (58.8% compared to 51.5%, *P* < 0.001).

Feed intake was affected by rumen and blood serum sample collections on days 6 and 14 with both groups showing a reduction, and so these data were removed from further analysis (Supp. Table 3). Overall, feed intake was significantly greater (*P* = 0.01) for wethers fed the pelleted diet compared with the loose hay diet; average dry matter intake increased by 17% (1.60 vs 1.86 kg/d, respectively) (Table 2). Similarly, average daily gain throughout the trial was greater (*P* = 0.0003) for wethers consuming the pelleted diet (0.24 +/- 0.02 kg/d) than for the loose hay diet (0.08 +/- 0.02 kg/d). Difference in ADG between pelleted and loose hay treatments was numerically greater (not specifically contrasted) after week 2 (*P* = 0.0003, 0.312 and 0.143 kg/d, respectively) than in the first week (*P* = 0.02, 0.169 and 0.026 kg/d, respectively). Although feed intake was increased in the pelleted diet, FCR was significantly (*P =* ≤ 0.02) improved in week 1 (0.079 vs 0.013), week 2 (0.196 vs 0.121), and overall (0.129 vs 0.053) for pelleted versus loose hay diets, respectively. Estimated apparent NDF and ADF digestion using lignin as an internal marker was greater (*P* ≤ 0.004) for sheep fed loose hay than for sheep fed the pelleted diet (Table 2). Oppositely, apparent CP digestibility was greater (*P* < 0.0001) in sheep fed the pelleted diet than the loose hay diet.

**Supplemental Table 3.** Daily intake by diet.

| Day | Hay (kg) | Pellet (kg) |
| --- | --- | --- |
| 0 | 1.90 | 2.29 |
| 1 | 1.72 | 1.93 |
| 2 | 1.55 | 1.63 |
| 3 | 1.76 | 1.78 |
| 4 | 1.26 | 1.87 |
| 5 | 1.57 | 1.89 |
| 6 | 1.22 | 0.65 |
| 7 | 3.17 | 2.78 |
| 8 | 1.61 | 2.05 |
| 9 | 1.47 | 2.01 |
| 10 | 1.78 | 2.03 |
| 11 | 1.45 | 2.12 |
| 12 | 1.17 | 2.05 |
| 13 | 0.66 | 0.95 |
| 14 | NA | NA |

**Table 2.** Weight, intake, and efficiency values for wethers on loose-hay diet and pelleted-hay alfalfa diets. Apparent digestibility was estimated using lignin as an internal marker.

|  | Loose Hay | Pelleted | SE | P-value |
| --- | --- | --- | --- | --- |
| Feed Intake (kg/d) | 1.6 | 1.86 | 0.06 | 0.01 |
| Average Daily Gain (kg/d) | 0.084 | 0.24 | 0.018 | 0.0003 |
| Week 1 | 0.026 | 0.169 | 0.036 | 0.02 |
| Week 2 | 0.143 | 0.312 | 0.019 | 0.0003 |
| Feed Conversion Ratio | 0.0527 | 0.129 | 0.008 | 0.0002 |
| Week 1 | 0.0128 | 0.0786 | 0.0167 | 0.02 |
| Week 2 | 0.121 | 0.196 | 0.014 | 0.006 |
| Apparent Digestibilities |  |  |  |  |
| App DMd | 35.6 | 40.7 | 2.2 | 0.12 |
| App CPd | 38.4 | 57.5 | 2.4 | <0.0001 |
| App NDFd | 34.9 | 26.7 | 1.8 | 0.004 |
| App ADFd | 38.4 | 30.5 | 0.8 | <0.0001 |

### Blood parameters

Serum values are provided in Supplemental Table 3. Blood creatinine was not different between treatments (*P* = 0.18) but CPK tended to be increased in the pelleted treatment by 135 units/L (Table 3; *P* = 0.06). Total protein showed a tendency to be greater in the pelleted diet (*P* = 0.08) but there were no differences in albumin nor globulin measurements to explain the variation (*P* ≥ 0.20). Many other standard panel measurements (AST, ALKP, GLU, and both total and direct bilirubin were unaffected by treatment (*P* ≥ 0.37). Blood urea nitrogen was greater in both treatment groups than the normal reference range for sheep [59] but pelleted treatment sheep were also greater than loose hay fed sheep (*P* = 0.02) by 4,25 mg/dL. Blood Ca was slightly greater in the pelleted treatment (*P* = 0.05) while P was decreased by 1.5 mg/dL in the pelleted diet (*P* = 0.0008). Finally, total blood CO_2_ was decreased within the pelleted treatment group but both treatment estimates are well within the range of normal for this test.

**Table 3.** Blood serum parameters for wethers fed either a loose alfalfa hay or pelleted alfalfa diet.

| Parameter<sup>1</sup> | Loose Hay | Pelleted | SE | *P*-value |
| --- | --- | --- | --- | --- |
| CPK | 299.49 | 434.64 | 43.06 | 0.06 |
| CREAT | 0.682 | 0.79 | 0.052 | 0.18 |
| AST | 113.99 | 114.73 | 7.93 | 0.95 |
| ALKP | 192.39 | 200.4 | 17.38 | 0.75 |
| T_PRO | 6.49 | 6.88 | 0.14 | 0.08 |
| ALB | 3.1 | 3.08 | 0.06 | 0.76 |
| GLOB | 3.27 | 3.51 | 0.12 | 0.2 |
| GLU | 63.62 | 66.68 | 2.43 | 0.4 |
| BUN | 27.7 | 31.95 | 1.08 | 0.02 |
| Ca | 10.29 | 10.52 | 0.07 | 0.05 |
| P | 7.48 | 5.97 | 0.2 | 0.0008 |
| Na | 143.92 | 142.92 | 0.36 | 0.09 |
| K | 4.7 | 4.82 | 0.12 | 0.48 |
| Cl | 104.41 | 104.66 | 0.34 | 0.61 |
| Mg | 2.27 | 2.27 | 0.05 | 0.97 |
| T_BILI | 0.406 | 0.42 | 0.031 | 0.75 |
| D_BILI | 0.07443 | 0.08721 | 0.009603 | 0.37 |
| CO<sub>2</sub> | 28.36 | 24.58 | 1.02 | 0.03 |
| HEM_Index | 0.278 | 0.234 | 0.048 | 0.54 |

<sup>1</sup>Blood parameter abbreviations are as follows: CPK: creatine phosphokinase, CREAT: creatinine, AST: aspartate aminotransferase, ALKP: alkaline phosphatase, T\_PRO: total protein, ALB: albumin, GLOB: globulin, GLU: glucose, BUN: blood urea nitrogen, T\_BILI: total bilirubin, D\_BILI: direct bilirubin, HEM Index: index of blood hemolysis.

### Bacterial richness in lamb rumen

Observed bacterial richness was very low in week 0, the end of the adaptation period (Fig. 2); it increased between days 0 and 7 (*P* < 2e-16), and days 0 and 14 (*P* < 2e-16), suggesting the rumen bacterial community was still in flux after two weeks of adaptation. Bacterial richness was significantly greater in the pelleted alfalfa diet at day 7 (*P* < 2e-16) and day 14 (P < 2e-16). Only day (F = 5.3686, *P* = 0.001) significantly explained overall bacterial community clustering in constrained ordination (Fig. 3); diet and diet x day were not significant. Overall, the constrained model was significant (F = 3.154, *P* = 0.001). However, day did not significantly contribute to variation in the community (varpart, variance = 0%), though community diversity contributed to 60% and day x diversity contributed to 39% of variance.

**Figure 2.**
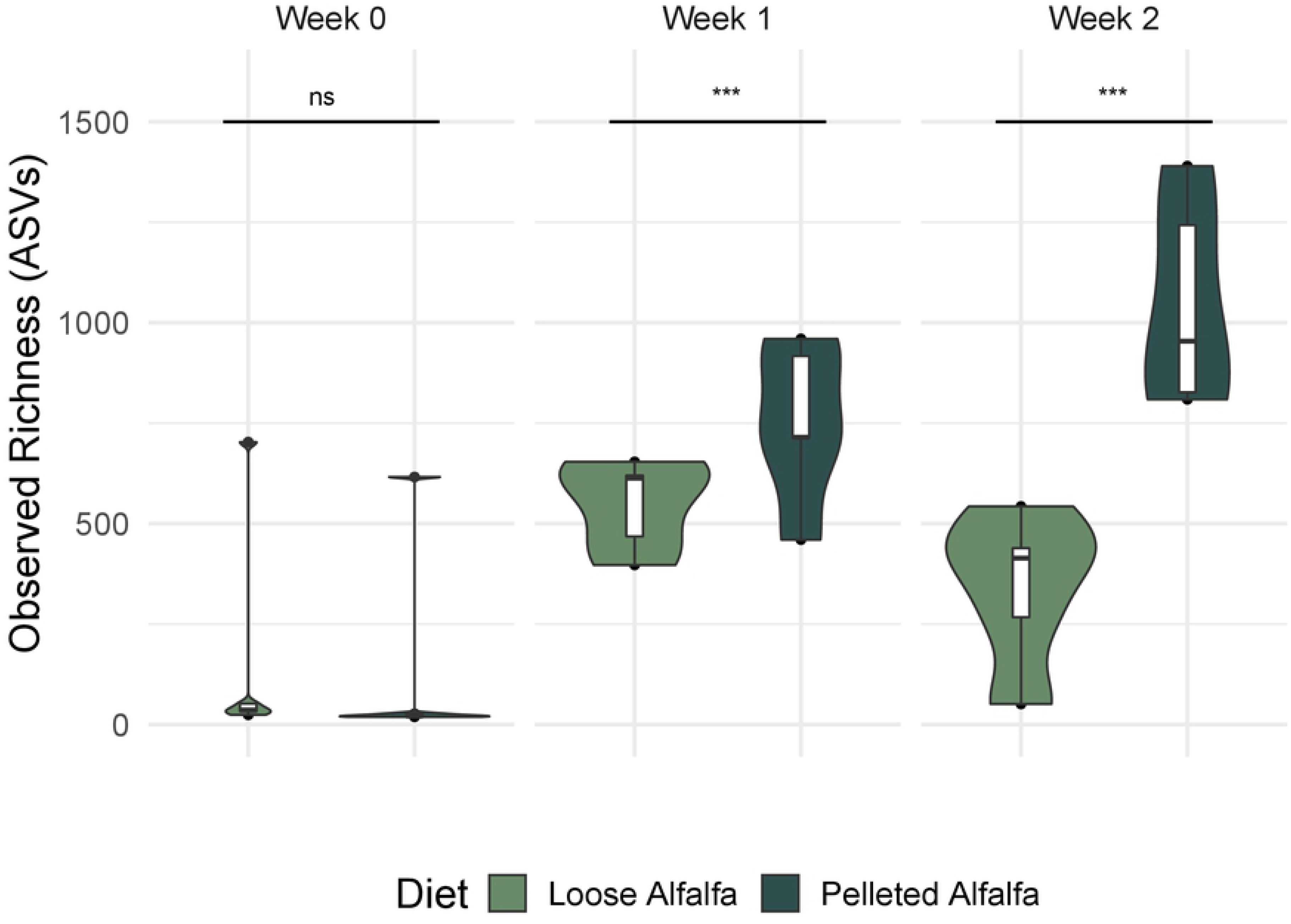
Observed bacterial richness in the rumen of wethers on loose-hay or pelleted-hay alfalfa diets. Significance was determined at *p* < 0.05 by linear mixed model with sheep ID as a fixed effect.

**Figure 3.**
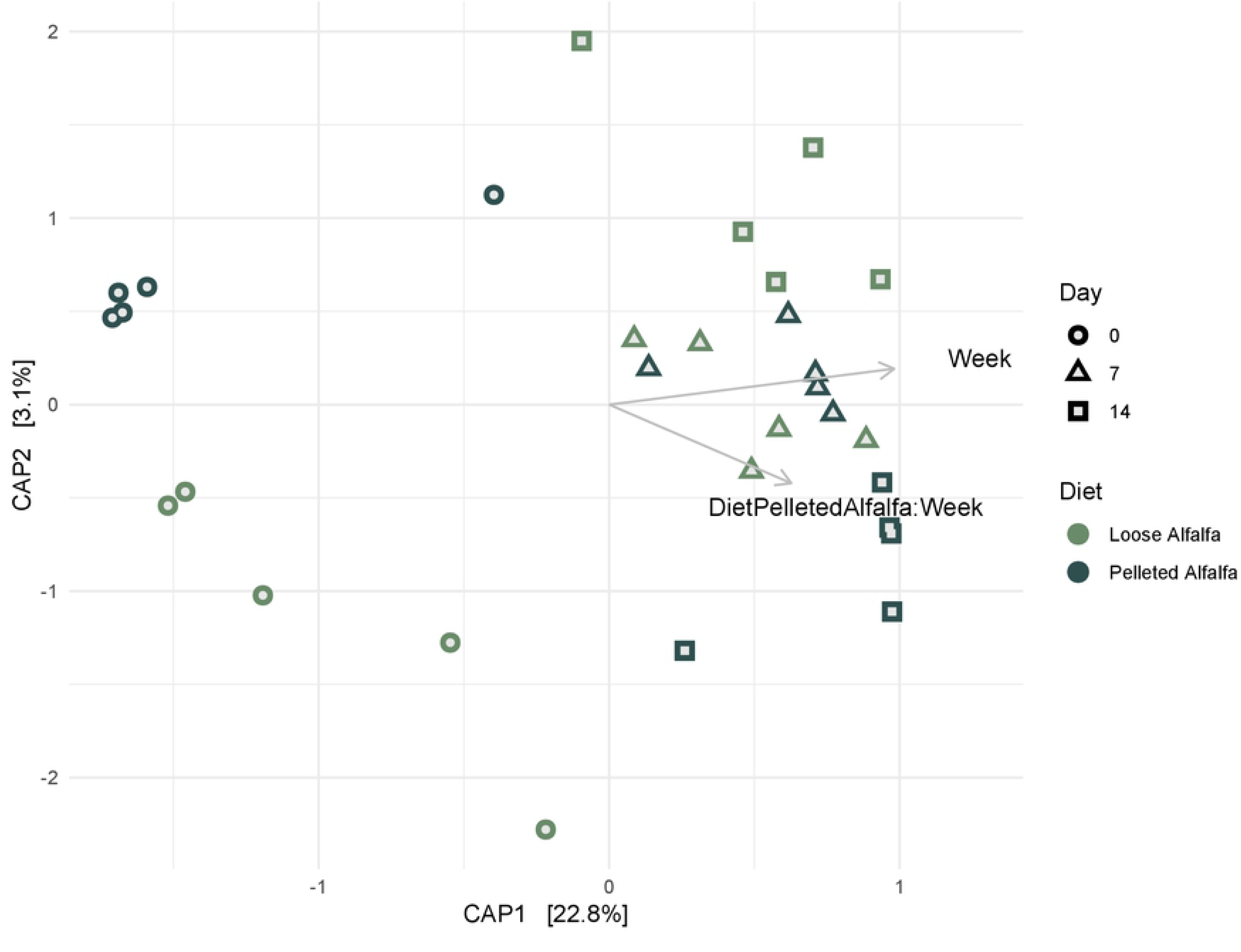
Distance-based redundancy analysis of Bray-Curtis Distance between rumen bacterial communities in wethers receiving loose-hay or pelleted-hay alfalfa diets.

Similarly, bacterial community composition clustered by sample day, followed by diet, in unconstrained ordinations (Fig. 4). Despite the increased richness, there was no significant diet:day interaction between bacterial communities based on unweighted (comparing samples based on OTU presence or absence) β-diversity (*P* = 0.31), however there was an effect of sample day (F = 2.44, R^2^ = 0.15, *P* = 0.0001) and dietary treatment overall (F = 1.50, R^2^ = 0.05, *P* = 0.037). Weighted (comparing samples based on OTU abundances) bacterial community distance, using Bray-Curtis Dissimilarity, was only significantly different by sample day (F = 4.64, R^2^ = 0.25, *P* = 0.0001).

**Figure 4.**
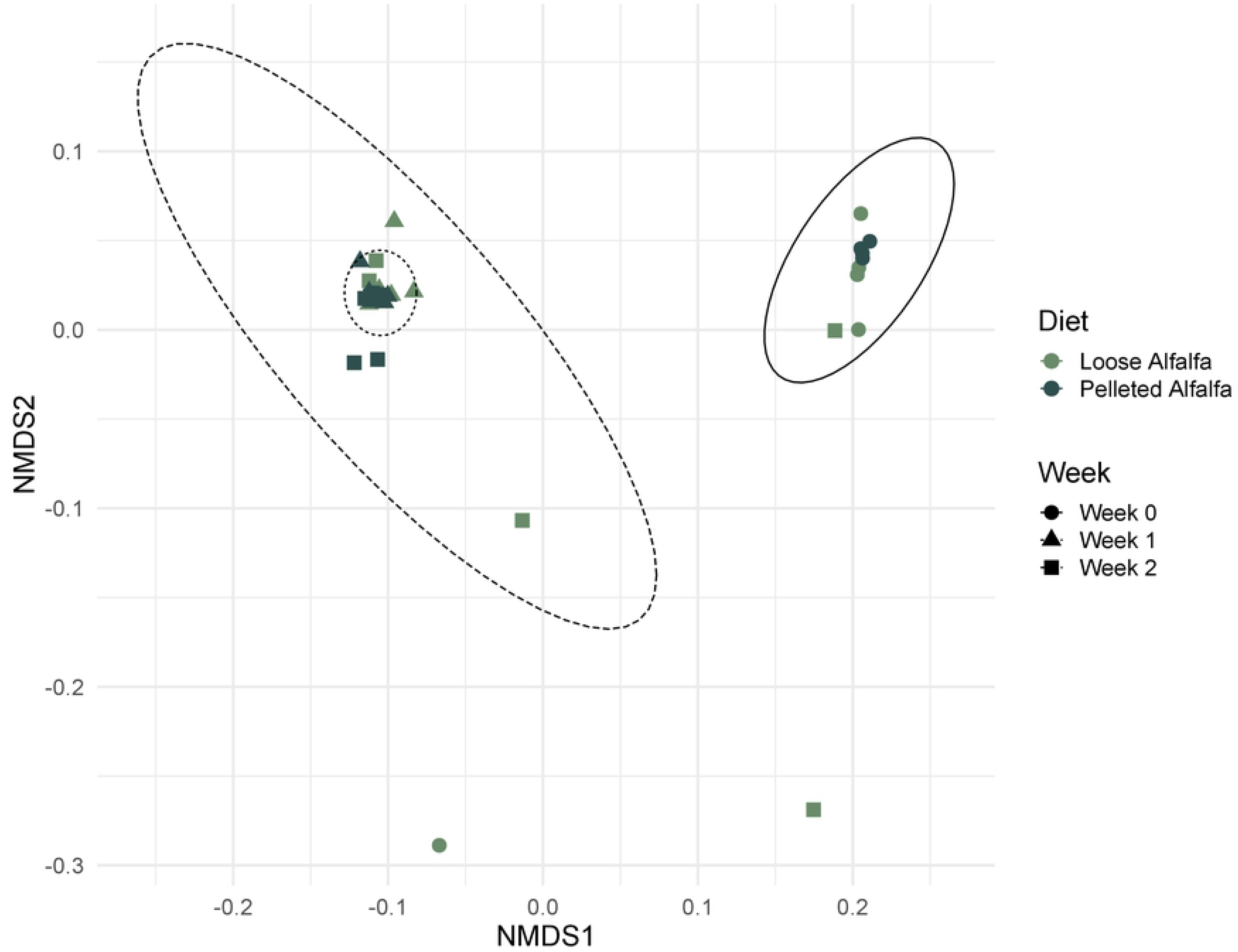
Non-metric multidimensional scaling (NMDS) plot of unweighted Jaccard distance between rumen bacterial communities in wethers receiving loose-hay or pelleted-hay alfalfa diets. Ellipses represent treatment weeks. Lowest stress = 0.0649, R^2^ = 0.9958.

When examining samples from day 7 and 14, to minimize any potential impact of dietary adaptation occurring between trial days 0 and 7 samples, 86 SVs were significantly discriminatory to the pelleted diet over loose-hay alfalfa diets, the 50 most abundant of which are presented in Figure 5. These included Ruminococcaceae UCG-005, *Succiniclasticum*, the RC9 gut group, and several *Prevotella*. Random forest prediction accuracy for diet was 80%, using only days 7 and 14 for reference, but was only 60% accurate in determining discriminatory communities using all sample days (0, 7, and 14; data not shown), despite the close clustering of samples observed in Figure 4. Likewise, just 30% accuracy was obtained in predicting bacterial communities for treatment x sample day (data not shown).

**Figure 5.**
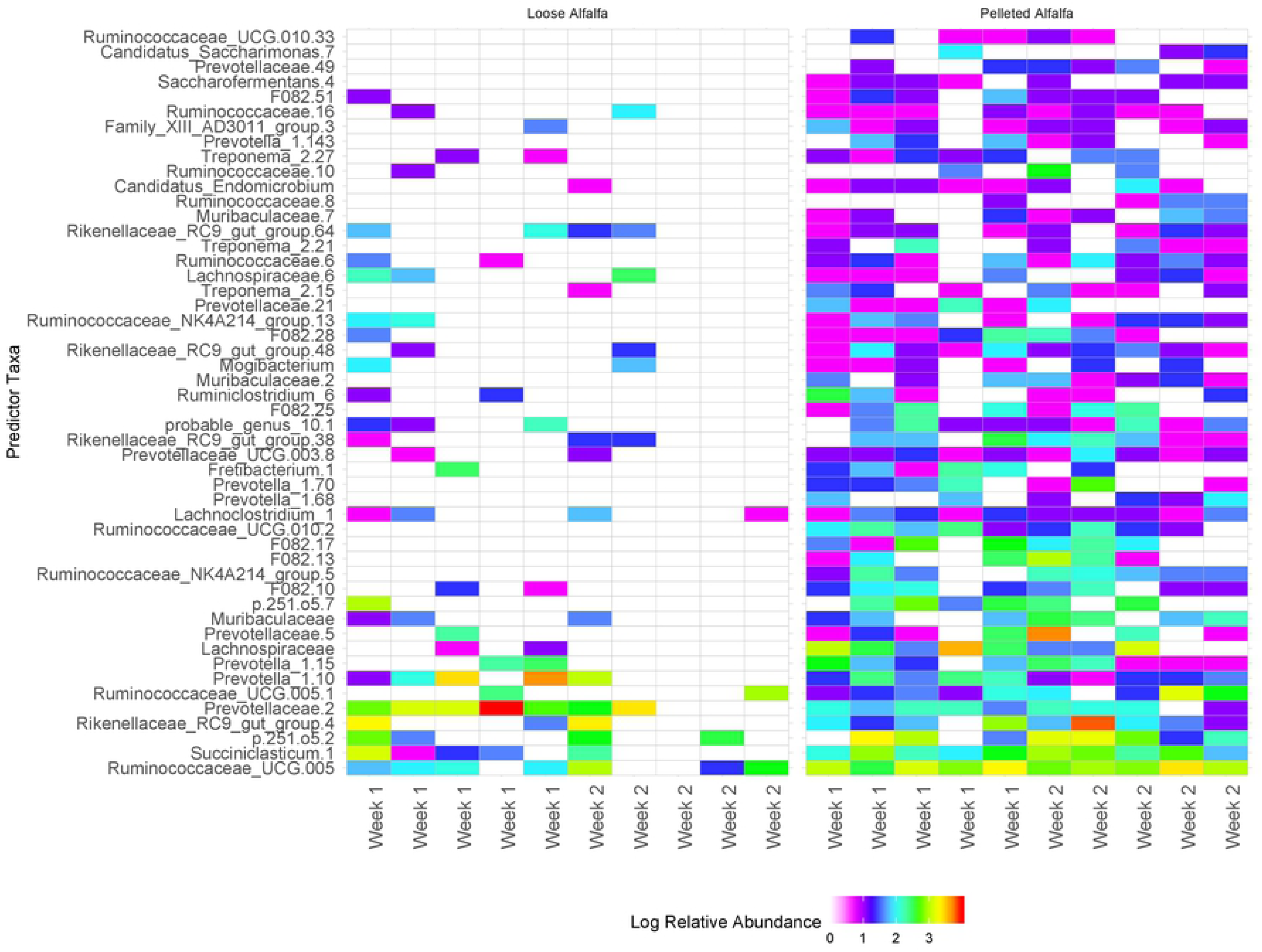
Discriminatory rumen bacterial sequence variance by treatment group for wethers receiving loose-hay or pelleted-hay alfalfa diet treatments. Significance (*p* < 0.05) determined by binomial test.

### Correlation of bacterial richness to performance parameters

The 86 rumen bacterial sequence variants (SVs) which were significantly discriminatory between lamb wethers on loose-hay and pelleted-hay alfalfa diets, as identified using random forest, were corelated against lamb weight and blood serum chemistry (Fig. 5). Spearman’s correlation revealed a number of bacterial taxa which were correlated with particular parameters. A number of the most abundant bacterial taxa identified in the rumen were significantly correlated with lamb growth and health, as well as with diet treatment (Figure 6). Body weight was positively correlated with several SVs in Prevotellaceae family, an SV in the Ruminococcaceae family, and an uncultured bacterium in the Bacteroidales order (p-251-05). The pelleted diet was positively associated with several *Succiniclasticum*, a Prevotella, and uncultured taxa in the Ruminococcaceae and Rickenellaceae families and Bacteroidales order. Several serum factors were correlated with lamb body weight (Table 3).

**Table 3.** Serum factors significantly correlated (Spearman’s) with weight in lambs.

| Serum Factor | R | p |
| --- | --- | --- |
| CL | 0.408037 | 0.025196 |
| CPK | -0.3864 | 0.034931 |
| CREAT | 0.435603 | 0.016125 |
| GLOB | 0.373562 | 0.04201 |
| T.BILI | -0.36713 | 0.045963 |
| T.PRO | 0.408708 | 0.024934 |

**Figure 6.**
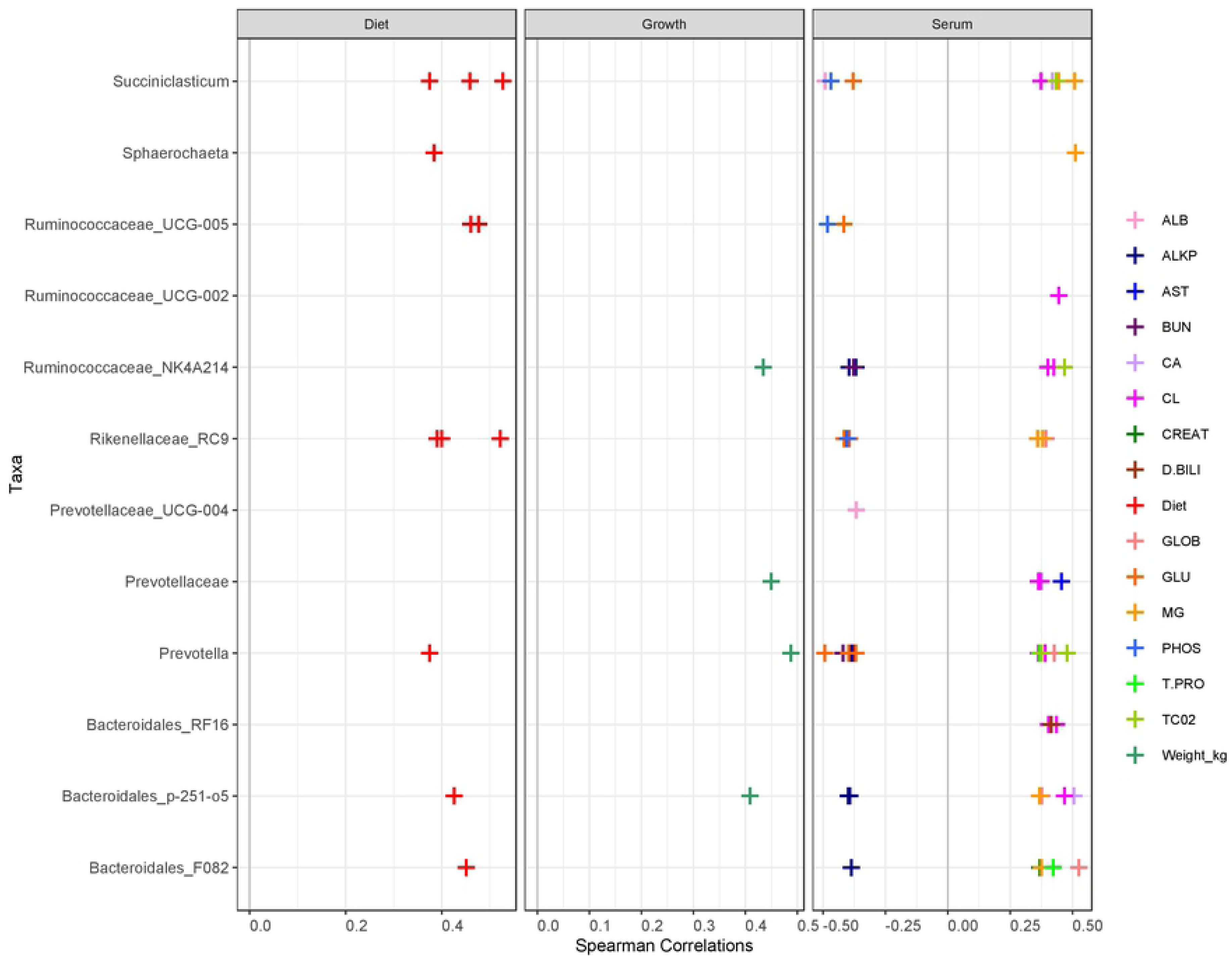
Correlations between the 50 most abundant rumen bacteria and lamb growth, serum parameters, and treatments. Only significant (*P* < 0.05) Spearman’s correlations are reported.

## Discussion

Despite the known effect of diet particle size on physical egress through the GIT, chemical accessibility and digestibility, and host absorption, the explicit effect of feed particle size has never been examined within the context of rumen microbial diversity. It is well established that forage-based diets promote increased bacterial diversity when compared to starch-based diets [60–63]. However, the role of particle size therein is unclear, as differences in the structural fiber composition and resulting bioavailability of nutrients that occur among plant species can select for distinct microbial communities [64]. In the present study, a smaller particle size feed increased bacterial diversity and sheep performance, likely by improving protein digestion of feed.

In other studies comparing the same diet across multiple particle sizes, no difference in average daily gain was observed by particle size regardless of an increase in DMI [42,43], which is in contrast to results presented here. Lambs fed the pelleted diet out-consumed and out-gained lambs fed the loose hay. Likely, increased DMI decreased digestibility for lambs fed the ground and pelleted diet [65]. This loss in digestibility was anticipated given a wealth of previous literature on fiber particle reduction, feed intake response, and reduced digestibility [66]. However, loss in digestibility was offset by net increase in digestible energy supply which combined with improved apparent CP digestibility contributed to improved gain in the pelleted diet lambs. The improvement in feed conversion ratio for pellet fed lambs is simply explained as a dilution of maintenance, where lambs eating and digesting more feed will have additional energy and protein in excess of requirement to allocate towards growth.

Some particle reduction studies using cows report loss of efficiency of microbial protein synthesis and total diet digestibility, perhaps in part due to a poor functioning rumen mat [67]. A smaller particle increases the available surface area for microorganisms to attach to feed but also increases passage rate from the rumen; the difference between ground and unground forage would be exacerbated in cows where particle reduction by mastication is less than in sheep [37]. A faster passage rate may sacrifice some ruminal fiber digestibility [68] yet increase microbial protein synthesis efficiency as bacteria compete to subsist in more rapid turnover environments [69,70]. In the present study, increased residue of NDF and ADF relative to lignin in the feces of sheep fed the pelleted diet indicate a poorer digestibility of fiber when provided in a ground, pelleted form. Studies have shown a U-shaped relationship of lignin content in the residual material of feces to particle size [22], supporting the idea that an overly-fine diet may not be beneficial. Given the effect of decreasing particle size on rumen retention time [22], this loss in fiber digestibility observed in the present study was expected and pairs with foundational work on the effect of grinding on fiber digestibility [71]. Furthermore, mechanical processing and pelletization of feed, while creating more surface area, can also compact the open space created by complex carbohydrate structure, thereby reducing the water-holding capacity of the feed and increasing its density, which may have contributed to lower microbial digestion [37].

In the present study, most serum values were within normal ranges, with the exception of creatine phosphokinase (CPK) at week 1 for pelleted-hay sheep, which was high for some individuals [72]. This spike in CPK presumably led to the detected significance for pelleted treatment to increase CPK compared with the loose hay diet. Traditionally, CPK has been associated with selenium deficiency induced white muscle disease in sheep, high sulfur-induced polioencephalomalacia [73,74], or copper toxicity [75]. However, none of these explain the current treatment difference and it is possible the greater CPK values may reflect short-term sampling stress [76]. Digestibility crates notoriously disrupt animal behavior; perhaps the strong flocking instinct of sheep created high stress levels for some individuals that was not reflected in feed intake data [59,76].

Blood urea nitrogen concentration in the present study was increased in the pelleted diet. The pelleted diet ground and thus more prone to greater (and estimated apparent) CP digestibility; increased BUN in the pelleted treatment supports this data as greater rumen degradable protein or (leading to ruminal ammonia concentration) in the diet will increase circulating urea recycling to the rumen [77]. The high concentration of BUN in both treatments indicates degradable protein was in excess for both treatments. More research needs to be done on the appropriate balance of degradable to undegradable protein for growing sheep in order to prevent systemic overfeeding of degradable protein in lambs.

Limited small ruminant studies do not report an effect of particle size per se on blood biochemical parameters in goats fed rice straw [78] or lambs fed alfalfa [35]; however, some studies do report an effect on some serum metabolites and protein digestion values by single bacterial species when introduced to gnotobiotic animals [79–82] A review of probiotic in lambs reported both increased and decreased levels of blood urea nitrogen, glucose, and creatine, citing changes in fermentation and ammonia cycling resulting from altered rumen bacterial diversity [83]. In the present study, blood urea nitrogen, calcium, creatine, globulin, magnesium, and total protein were positively correlated many bacterial taxa, while sodium, phosphorous, potassium, and total carbon dioxide were negatively correlated with many other bacteria. Nearly all bacteria were positively correlated with weight, except for *Fretibacterium*, previously identified in the human oral cavity [84]; it does not produce acid from sugars but which can synthesize acetic acid and propionic acid [84]. Perhaps a microbial adaptation to the particle size treatment is partially responsible for minor shifts in blood serum measurements in the present study.

Observed bacterial richness was very low in week 0 at the end of the adaptation period, and then increased for the rest of the study, suggesting the rumen bacterial community was still in flux even after two weeks of adaptation. Similarly, samples clustered largely by week, indicating the need for longer dietary adaptation periods in ruminant studies. The pelleted-hay diet recruited significantly more bacterial richness, including common rumen inhabitants known to ferment a variety of carbohydrates, such as *Prevotella, Ruminococcus, Succiniclasticum*, and members of the RC9 gut group. Yet despite the increase in average daily gain and feed efficiency, there was no difference in residual fecal nutrients by diet except protein, which was reduced in pelleted-hay lambs. It is possible that the improvement in animal performance was not from fiber digestion but from enhanced access to microbial and dietary protein resulting from a combination of greater increased microbial growth due to improved access to dietary nutrients making available more surplus energy for increasing microbial biomass, and an increased flow rate of digesta and associated microbiota from the rumen.

## Acknowledgements

This study was generously supported by the Montana Agricultural Experiment Station (project MONB00113) and is a contributing project to the Multistate Research Project, W4177, Enhancing the Competitiveness of U.S. Beef (MONB00195).

